# Voice identity invariance by anterior temporal lobe neurons

**DOI:** 10.1101/2024.12.19.629267

**Authors:** M. Giamundo, R. Trapeau, E. Thoret, L. Renaud, T. G. Brochier, P. Belin

## Abstract

The ability to recognize speakers by their voice despite acoustical variation plays a significant role in primate social interactions. While neurons in the macaque anterior temporal lobe (ATL) show invariance to face viewpoint, whether they also encode abstract representations of caller identity is not known. Here we show that neurons in the voice-selective ATL of two macaques show invariance to voice identity via dynamic representations that minimize within-caller neuronal distances while maintaining distinct neural trajectories for different individuals. A small proportion of highly identity-selective neurons plays a central role although less selective neurons are also informative. Our findings provide a neural basis for voice identity recognition in primates and highlight the ATL as a key hub for integrating perceptual voice features into higher-level identity representations.

## Introduction

Human listeners readily extract speaker identity information from even brief voice samples—an ability at the core of our social interactions. Non-human primates share a similar ability as shown by both field and laboratory studies (Zoloth & Green, 1979; Cheney & Seyfart, 1980; Gouzoules et al., 1984; Hauser, 1991). Although neuroimaging studies start shedding light on the neural circuits involved in voice processing by humans, an understanding of how identity information is encoded at the level of individual neurons is still lacking, in contrast to face identity processing (Freiwald & Tsao, 2010; Chang & Tsao, 2017; Landi et al., 2021; She et al., 2024).

Numerous studies in humans and non-human primates suggest that a key biological substrate for identity processing is the ATL as a convergence site for different sensory streams (for reviews, see Olson et al., 2013; Perrodin et al., 2015; Gainotti, 2018; Deen et al., 2023). In the auditory modality, the anterior Temporal Voice Area (aTVA) is an ATL region specialized in the processing of conspecific vocalizations in humans and macaques (Petkov et al., 2008; Perrodin et al., 2011; Pernet et al., 2015; Bodin et al., 2021; Giamundo et al., 2024).

Functional MRI studies found that the aTVA shows adaptation to speaker identity in both humans (Belin & Zatorre, 2003) and macaques (Petkov et al., 2008), i.e. decreases its neural activity when different consecutive stimuli are from a same speaker, suggesting abstract representation of speaker identity (Kriegeskorte et al., 2007; Formisano et al., 2008; Andics et al., 2010). How such an abstract, variability-robust representation of speakers emerges at the level of individual neurons is still largely unknown. A single study so far has investigated the processing of caller identity in macaque aTVA neurons, showing that a small subset of these neurons differentiates between individuals more than between call type (Perrodin et al., 2014), but whether and how they are involved in caller recognition is unclear.

Here we investigate to what extent neuronal population dynamics in the macaque aTVA process voice identity information. As suggested by electrophysiological studies of face identity processing (Chang and Tsao, 2017; Landi et al., 2021), we hypothesize that individual voice identities are abstractly encoded by neurons that recognize an individual despite acoustical variation. Given the existence of ‘grandmother’ neurons selective to a single identity in the hippocampus (Quiroga et al., 2005), we also examine whether and how identity-selectivity contributes to caller recognition.

To test these hypotheses, we used fMRI-guided electrophysiology in two macaques to examine the activity of their aTVA neurons during exposure to vocalizations from five different macaque callers. Our findings reveal that (1) aTVA population of neurons almost perfectly encodes voices by identity, (2) they are particularly involved in recognition by minimizing neural distances between different calls from a same caller, and (3) voice identity recognition mainly relies on neurons with high selectivity for specific identities although mildly identity-selective neurons were also informative.

## Results

### aTVA neurons exhibit strong voice identity selectivity

To investigate the neural bases of voice identity processing, we analyzed spiking activity from single neurons of two female rhesus monkeys, M1 and M2, implanted with chronic multi-electrode arrays in the fMRI-localized aTVA (Fig. 1A; Bodin et al., 2021; Giamundo et al., 2024). The electrophysiological recordings took place while the monkeys performed active detection of a pure tone interspersed amongst a set of complex vocal stimuli (Fig. 1B). The set of stimuli consisted of 25 natural vocalizations: 5 different “coo calls” from each of 5 macaque individuals (Fig. 1C). Coo calls, also termed “contact calls”, are harmonically rich affiliative vocalizations commonly produced by macaque species (see Hauser, 1991 for a review). We chose to focus on this class of vocalizations due to macaques’ demonstrated ability to recognize individual identities based on coos (Rendall et al., 1996). The 5 monkeys that produced the vocalizations were unknown to our two monkeys.

**Fig. 1.**
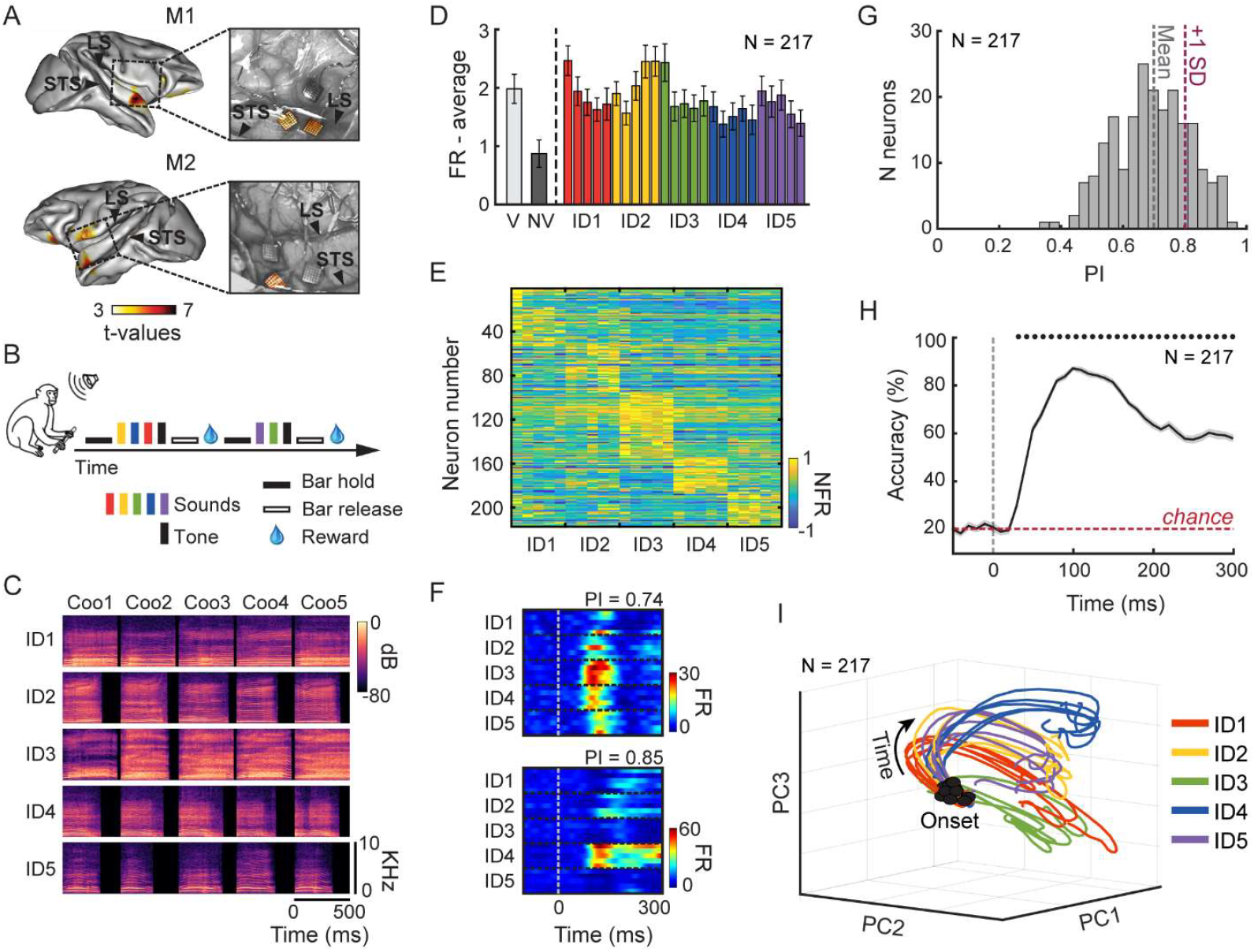
aTVA neuronal activity is modulated by voice identity. (A) Implantation sites of high-density chronic electrode Utah Arrays in monkeys M1 and M2. Color scale indicates t-values of fMRI contrast between conspecific macaque vocalizations vs. non-vocal sounds. Utah arrays analyzed here (highlighted in the pictures of cortical surface during surgery, right insets) were implanted in cortical areas close to the fMRI-identified aTVA peaks. STS: Superior Temporal Sulcus. LS: Lateral Sulcus. (B) Pure tone detection task. Monkeys were trained to release a bar when a pure tone was randomly presented amongst other stimuli to obtain the reward. (C) Spectrograms of the main stimulus set used in the experiment, comprising 25 natural ‘coo’ calls produced by five individual macaques (identities, IDs). (D) Average population response (mean ± SE) to macaque vocalizations (V) and non-vocal sounds (NV) of the localizer stimulus set, and to the 25 voice identity (ID) stimuli, with the color corresponding to each identity. (E) Neurons responsiveness to the 25 voice identity stimuli. Each row represents the average response of a single neuron to each stimulus, calculated over the 200 ms window following stimulus onset and normalized to its absolute maximum response across stimuli (see Methods). Neurons are sorted based on the identity eliciting their strongest response. (F) Mean response time courses of two typical neurons to the 25 voice identity stimuli. The first 5 rows are responses to the 5 coo calls of individual 1 (ID1), and so on. Responses are aligned to the stimulus onset. PI: Preference Index. (G) Histogram of Preference Indices (PIs), reflecting each neuron’s selectivity for a single identity during the 200 ms window following stimulus onset. A PI of 0 indicates equal responsiveness across all five identities, while a PI of 1 reflects exclusive responsiveness to a single identity. Vertical dashed lines represent the mean PI (Mean) and the mean plus one standard deviation (+1 SD). Note the large proportion of neurons with a high PI values. (H) Time-resolved classification accuracy (mean ± SE) of linear classifier trained to discriminate between identities (chance level: 20%) based on the population spiking activity. Black dots above indicate time bins of significantly above-chance classification accuracy (permutation tests, p < 0.0004; see Methods). (I) Neural trajectories of the responses to the 25 voice identity stimuli in the state space defined by the first three PCs, from - 100 to +300 ms relative to stimulus onset. Trajectories are colored-coded by identity.

We focused our analyses on 217 well-isolated auditory-responsive neurons (173 neurons from M1 and 44 neurons from M2), i.e. neurons responding to at least one of the 25 stimuli (see Methods). Below, we combine the data from the two monkeys because we did not find any marked differences between individuals, but see Figs. S1-S3 for the main results separately for each animal.

We initially assessed the selectivity of these neurons to voices by comparing their spiking activity in response to the 25 voice identity stimuli with their responses to a localizer consisting of 12 macaque vocalizations and 12 non-vocal sounds presented at the beginning of each recording session (see Methods). The aTVA neuronal population exhibited a stronger response to macaque vocalizations compared to non-vocal sounds (Figs. 1D, S1A,F). Furthermore, Voice Selectivity Indices (VSIs) contrasting, for each neuron, firing rates of the 12 macaque vocalizations and the 12 non-vocal sounds of the localizer (see Methods), showed more selectivity for macaque vocalizations than for non-vocal sounds (42% of neurons with a VSI > 0.33 – i.e. neuronal response to voices at least two times stronger than to non-vocal sounds – compared to 27% of neurons with a VSI < -0.33), confirming the voice sensitive nature of this region.

Although the average spiking activity of the aTVA neuronal population did not show substantial differences across the five macaque identities (Figs. 1D, S1B), individual neurons exhibited diverse response patterns, ranging from broad responses to multiple identities to highly specific responses for a single identity (Figs. 1E-F, S1C). We quantified the selectivity of neurons to one identity using a preference index (PI). The PI reflects a neuron’s ability to distinguish between identities, with values near 1 indicating strong selectivity for one specific identity, and values near 0 indicating similar responses to all five identities (see Methods). Overall, aTVA neurons exhibited strong identity selectivity with a mean PI of 0.7 (± 0.1 SD), suggesting the involvement of this region in encoding voice identity (Figs. 1G, S1D-E). No correlation was observed between the PIs and VSIs of neurons (Pearson correlation, rho = 0.09, p = 0.18).

### Voice identity is encoded at the population level

We investigated the role of the aTVA neuronal population in encoding voice identity by training a machine-learning classifier (maximum correlation coefficient) to distinguish between identities based on spiking activity (20 ms time windows every 10 ms steps). Decoding accuracy was significantly above chance (chance = 20%) from 30 ms after stimulus onset and persisted throughout the entire duration of the sound (all p’s < 0.0004, permutation tests), reaching 88% of accuracy at 100 ms after onset (Fig. 1H). The classifier performed consistently across all identities, with no significant differences in misclassification rates (Fig. S2B, top panel; refer to figs. S2A-B for decoding results specific to each monkey).

We conducted the same decoding analysis on voice-selective and non-vocal selective neurons to explore whether voice selectivity affects the ability to classify between identities (Fig. S2C). Both subpopulations exhibited classification accuracy significantly above chance from 40 ms after onset (all p’s < 0.0004, permutation tests), although accuracy was consistently higher for voice-selective neurons during multiple time windows (40-120 ms, 140-200 ms, and 240-300 ms; all η values < 0.05; see Methods).

To visualize the dynamics of the neuronal population’s responses, we applied a dimensionality reduction approach by principal component analysis (PCA). This allowed us to represent the responses to the 25 vocalizations as neural trajectories evolving in a low-dimensional state space. Fig. 1I shows the neural trajectories across the stimulus presentation time, plotted in the subspace defined by the first three principal components (explained variance: PC1 = 15%, PC2 = 13%, PC3 = 8%). Following stimulus onset, trajectories rapidly converged for vocalizations from the same identity and diverged for different identities in the state space (Fig. 1I). To quantify this pattern, we computed the Euclidean distances between neural trajectories, revealing a significant increase in the distance of trajectories between identities compared to the distance of trajectories within each identity from stimulus onset (from -4 ms throughout the stimulus presentation time, all t(298) > 3.7905 and all p’s below Bonferroni-corrected p-value = 0.0001). These findings show that the aTVA population robustly encodes voice identity through dynamic and distinct neural patterns.

### aTVA neurons prioritize identity Recognition over Discrimination

Voice identity processing in humans has been proposed to consist of two main abilities (Van Lancker & Kreiman, 1986; Lavan et al., 2019; Schweinberger et al, 2014): *1) Voice Discrimination*: perceiving that two voice samples are from separate individuals (‘telling voices apart’), primarily based on low-level acoustical differences; and *2) Voice Recognition*: perceiving that two voice samples have been produced by a same individual (‘telling voices together’), a higher-level ability requiring abstraction from within-speaker acoustical variability. Unraveling how these dual processes occur in populations of individual neurons is crucial for understanding the neuronal mechanisms underlying voice identity processing and more generally, person perception.

We sought to disentangle the neural patterns underlying these two key components of voice identity processing by applying Representational Similarity Analysis (RSA; Kriegeskorte et al., 2008). This technique enables direct comparisons between brain activity representational patterns and theoretical models. We computed a 25×25 Neuronal Representational Dissimilarity Matrix (RDM) every 10 ms time bins during stimulus presentation. Neuronal RDMs captured the dissimilarity in population spiking activity – measured as Euclidean distance in multi-neuron space –for each pair of the 25 voices, using a 50 ms window centered on the bin (Fig. 2A, left panels; Methods).

**Fig. 2.**
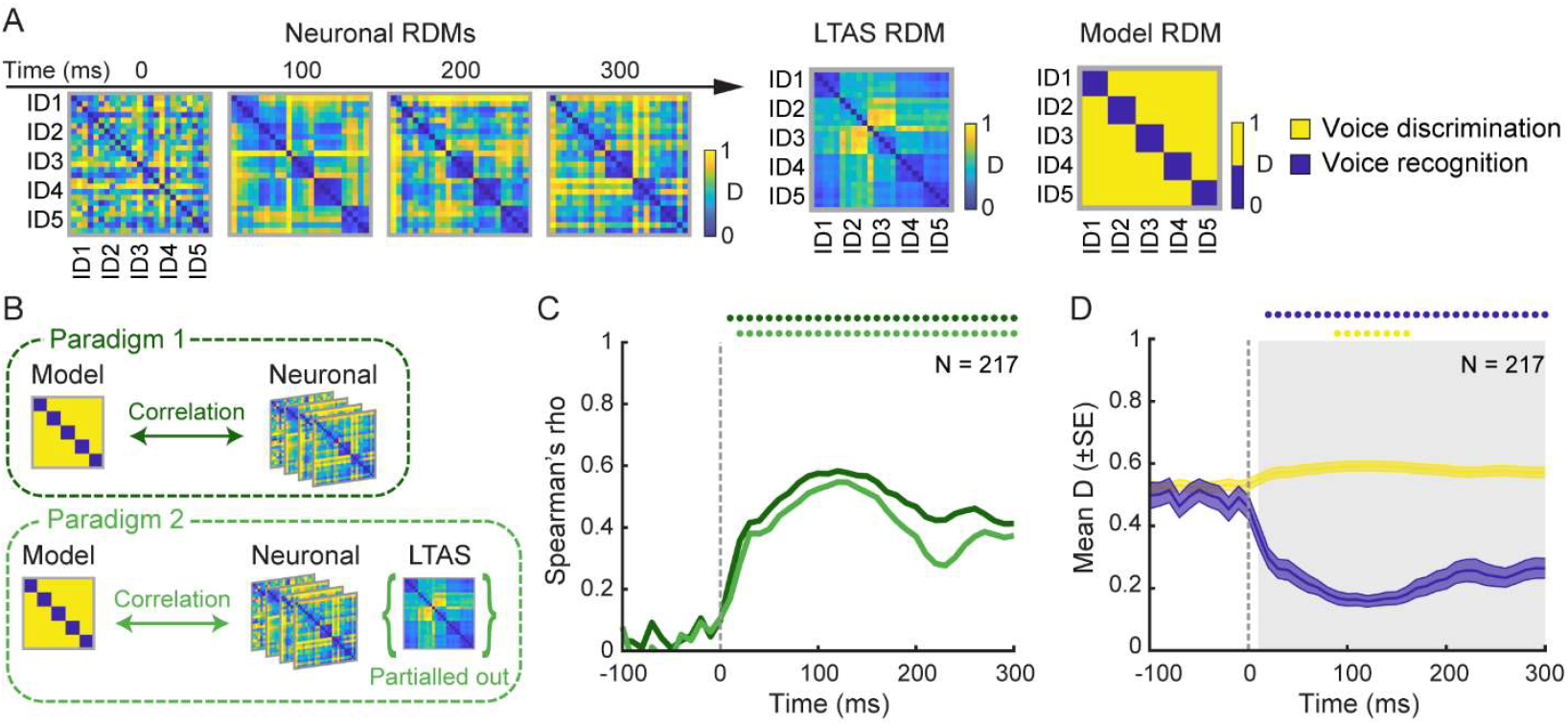
Neural patterns underling voice discrimination and voice recognition processes. (A) The different RDMs compared. Left panels: Neuronal RDMs capturing spiking differences across all pairs of the 25 voice identity stimuli, as a function of stimulus presentation time. Color scale indicates normalized pairwise distance rankings. Middle panel: long-term average spectrum (LTAS) RDM. Right panel: theoretical Model RDM representing ideal distinction of voice discrimination and voice recognition patterns. D: dissimilarity. (B) The two analysis paradigms used to test the association of Neuronal RDMs with the Model RDM, taking into account the contribution of low-level features of the stimuli by the LTAS RDM. (C) Time course of correlations (Spearman) derived from Paradigm 1 (dark green curve) and Paradigm 2 (light green curve). Colored dots above indicate time bins of significant correlation (Bonferroni-corrected p < 0.0012). (D) Dissimilarity values (mean ± SE) of the portions of Neuronal RDMs corresponding to voice discrimination (yellow curve) and voice recognition (blue curve) processes, predicted by the theoretical model. The gray shaded area indicates time bins of statistically significant association with the theoretical model (bootstrapped two-sample t-tests, Bonferroni-corrected p = 0.0012). Colored dots above indicate time bins with a significant difference with the baseline period (paired t-tests, Bonferroni-corrected p < 0.0012).

We compared the time-resolved Neuronal RDMs with a theoretical model designed to capture both voice Discrimination and Recognition processes (Fig. 2A, right panel). The model depicts an ideal pattern of dissimilarities between pairs of stimuli, where voices from the same identity are grouped together, showing no dissimilarity in their neural responses (Recognition, shown in blue), while voices from different individuals exhibit maximum dissimilarity (Discrimination, shown in yellow).

To account for the potential influence of low-level acoustic features on neuronal responses, we also computed an RDM based on the long-term average spectrum (LTAS) of the vocalizations (Fig. 2A, middle panel). Indeed, pairs of vocalizations within the same individual could have more similar low-level acoustical features than across different individuals. This measure reflects differences in spectral shape, encompassing both the source and filtering characteristics of each sound (Ghazanfar & Rendall, 2008).

To test the alignment of neuronal activity with the theoretical model, we used two analysis paradigms (Fig. 2B; Lavan et al., 2024). In Paradigm 1, we directly correlated time-resolved Neuronal RDMs with the theoretical Model RDM. In Paradigm 2, we repeated the analysis while partialling out the LTAS RDM to isolate contributions from higher-level features.

In Paradigm 1, Spearman’s rank correlations between Neuronal and Model RDMs were strong, emerging at 10 ms post-stimulus onset and persisting throughout stimulus presentation (Bonferroni-corrected p < 0.0012; Fig. 2C). When controlling for the LTAS RDM in Paradigm 2, correlations remained robust, albeit slightly reduced, from 20 ms post-stimulus onset (Fig. 2C; Bonferroni-corrected p < 0.0012). These findings suggest that although LTAS contributes marginally to aTVA computations, higher-level features are the primary drivers of voice identity representation.

Next we directly compared Discrimination to Recognition by isolating the portions of the Neuronal RDMs corresponding to these processes. We observed that the association with the theoretical model reached significance at 10 ms post-stimulus onset and persisted throughout sound presentation (bootstrapped two-sample t-tests, Bonferroni-corrected p < 0.0012; Fig. 2D). Moreover, neural distances between voices of different identities (Discrimination) showed only a modest increase compared to the baseline period (from -100 ms to stimulus onset; Fig. 2D), becoming significant between 90-160 ms (paired t-tests, Bonferroni-corrected p < 0.0012). In contrast, neural distances within the same identity (Recognition) decreased significantly from 20 ms post-stimulus onset, with this effect persisting throughout the stimulus duration (paired t-tests, Bonferroni-corrected p < 0.0012).

These results indicate that the aTVA population predominantly supports the Recognition and grouping of variations in a single individual’s voice. The limited influence of low-level acoustics further emphasizes the role of higher-level processing in driving these computations, suggesting that aTVA neurons prioritize identity recognition over simple acoustical discrimination.

### Identity-selective neurons are important but not essential to voice identity processing

We observed that many neurons exhibited strong selectivity for specific individuals (Figs. 1E-F, 3A), raising the question of whether voice identity information is distributed across the entire population, requiring all neurons for representation, or if a smaller subset of identity-selective neurons contains sufficient information to represent individuals.

To address this, we compared the contributions of identity-selective neurons to population-level processing with those of more broadly tuned neurons. We classified neurons with preference index (PI) values greater than one standard deviation above the mean (PI > 0.8) as identity-selective (N = 44; Fig. 3B). These neurons varied in their preferred identity, determined by their tuning profiles, and were differently distributed across the recording arrays (Fig. S3A). For comparison, we selected a subsample of the same size (N = 44) with the lowest PI values, representing non-identity-selective neurons (mean PI ± SD = 0.5 ± 0.05; Fig. 3B).

**Fig. 3.**
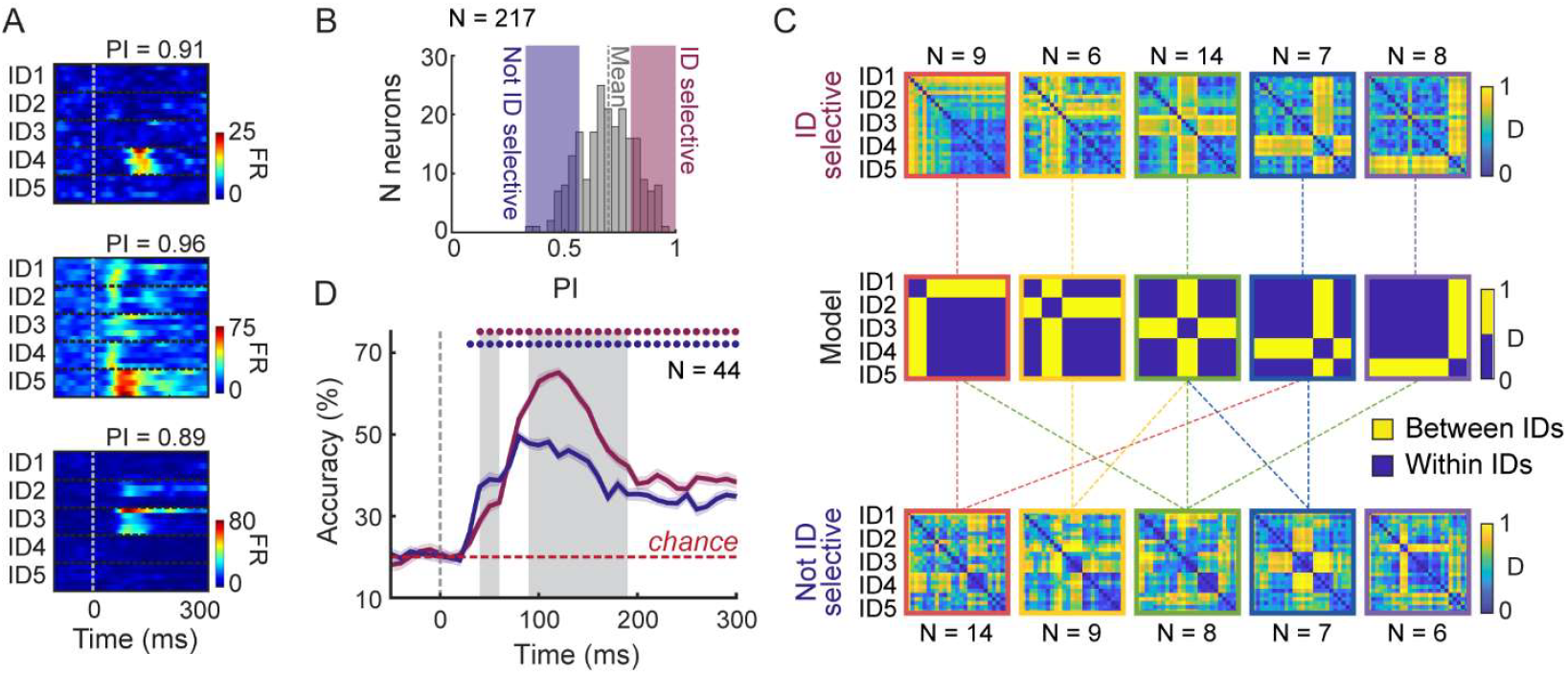
Contribution of identity-selective neurons to voice identity processing. (A) Mean response time courses of three typical identity-selective neurons to the 25 voice identity stimuli. PI: Preference Index. (B) Histogram of Preference Indices (PIs) as in Fig. 1G, highlighting n=44 identity-selective neurons (ID selective; in pink) and n=44 non-identity-selective neurons (Not ID selective; in violet). (C) Top and bottom panels: Neuronal RDMs computed from identity-selective (top) and non-identity-selective (bottom) neurons grouped by their preferred identities. Color scale indicates normalized pairwise distance rankings. Middle panels: theoretical Model RDMs representing ideal distinction of a single identity from all others. Dashed lines indicate significant comparisons between Neuronal and Model RDMs (bootstrapped two-sample t-tests, Bonferroni-corrected p < 0.001). (D) Accuracy (mean ± SE) in the classification between identities (chance level: 20%) for identity-selective (pink curve) and non-identity-selective (violet curve) neurons. Colored dots indicate time bins with significantly above-chance classification accuracy (permutation tests, p < 0.0004). The gray shaded areas indicate time bins with significant difference between classification accuracies of identity-selective vs. non-identity-selective neurons (η < 0.05).

Figure 3C shows Neuronal RDMs for identity-selective (top panels) and non-identity-selective (bottom panels) groups of neurons (defined by their preferred identity based on tuning profiles). RDMs from identity-selective groups of neurons closely matched (bootstrapped two-sample t-tests, Bonferroni-corrected p < 0.001) corresponding theoretical models representing ideal patterns in which a single identity is separated from all others (middle panels). In contrast, RDMs from non-identity-selective groups of neurons exhibited weaker or inconsistent alignment with these theoretical models (bootstrapped two-sample t-tests, Bonferroni-corrected p < 0.001; Fig. 3C).

We then assessed how identity-selective and non-identity-selective subpopulations encode voice identity using the same decoding approach applied to the full neuronal population (Fig. 1H). Both subpopulations classified between identities above chance (30 ms post-stimulus onset for non-identity-selective neurons and 40 ms for identity-selective neurons, all p’s < 0.0004, permutation tests; Fig. 3D). However, their performance differed in latency and strength (η values < 0.05; see Methods). Identity-selective neurons achieved higher classification accuracy, peaking at 120 ms post-stimulus onset, albeit with a slightly delayed onset. Conversely, non-identity-selective neurons demonstrated faster but less accurate classification, peaking at 80 ms post-stimulus onset (Fig. 3D). These results suggest that identity-selective neurons encode more detailed voice identity information in their spiking activity compared to broadly tuned neurons.

Despite these differences, both subpopulations exhibited similar temporal dynamics in coding stability. In both subpopulations, the highest decoding accuracy was achieved when training and testing data were from the same time, indicating that the coding patterns for voice identity evolved dynamically over time (Fig. S3B-C). Additionally, there were no differences between the subpopulations in representing voice recognition and discrimination processes (Fig. S3D).

These findings suggest that while identity-selective neurons contribute prominently to recognizing specific individuals, effective voice identity processing relies on the distributed responses of the entire neuronal population, emphasizing the importance of both specialized and broadly tuned neural contributions.

## Discussion

Together, our results show that neurons in the voice-selective ATL are highly sensitive to caller identity and show invariant coding of individuals across varying acoustic inputs (Lavan et al., 2019) via rapid convergence of firing patterns across stimuli from the same caller. Conversely, the lack of marked increase in between-speaker distances indicates that low-level acoustic discrimination likely occurs at earlier stages of auditory cortical processing as suggested by clinical studies (Van Lancker & Kreiman, 1986). The strong correlation between neuronal and theoretical dissimilarity matrices, even after controlling for the long-term average spectrum, underscores the abstraction of acoustical variability into stable identity representations. The exact nature of these higher-order features is important matter for future investigations.

Such caller invariance parallels the viewpoint-invariant responses observed in face-selective neurons. Neurons of the most anterior ‘face patch’ AM - located within the ATL - fire similarly to images of individual faces despite variations in viewpoints, a phenomenon not observed in more posterior, earlier-stage face patches (Freiwald and Tsao, 2010). This parallel confirms similarities in the functional organization of face and voice processing (Belin et al, 2004), suggesting common neural coding strategies for addressing the computational problem of abstraction from low-level features independently of sensory modality (Yovel & Belin, 2013; Zhang et al., 2021).

We find that voice identity processing relies on a hybrid combination of both sparse and more broadly identity-tuned neurons, emphasizing the distributed nature of robust identity encoding. About a fifth of the neurons were strongly identity-selective and supported the highest classification accuracy, likely playing a pivotal role in precise identity recognition. A key question for future research is whether these highly selective neurons also participate in cross-modal identity representations, similar to the “concept cells” identified in the hippocampus (Quiroga, 2012; Tyree et al., 2023), integrating sensory information into abstract identity constructs.

These results align with previous studies in humans and macaques, which consistently underscore the ATL’s critical role in in processing high-level cognitive features related to identity (Olson et al., 2013; Perrodin et al., 2015; Landi et al., 2017). The anterior voice patch may represent the final processing stage for perceptual voice features before relaying this information to higher-level regions for potential integration with other identity signals, facilitating the formation of new associations (Eifuku et al., 2010) and the feeling of familiarity (Deen et al., 2024).

## Funding

Fondation pour la Recherche Medicale AJE201214.

Agence Nationale de la Recherche grants ANR-16-CE37-0011-01 (PRIMAVOICE), ANR-16-CONV-0002 (Institute for Language, Communication and the Brain) and ANR-11-LABX-0036 (Brain and Language Research Institute).

Excellence Initiative of Aix-Marseille University (A*MIDEX).

European Research Council (ERC) under the European Union’s Horizon 2020 research and innovation program (grant agreement no. 788240).

## Author contributions

Conceptualization: MG, RT, PB

Methodology: MG, RT, TGB, PB

Surgical procedure: LR

Investigation: MG

Formal analysis: MG

Visualization: MG, PB

Funding acquisition: MG, PB

Project administration: MG, PB

Supervision: PB

Writing – original draft: MG, PB

Writing – review & editing: MG, RT, ET, LR, TGB, PB

## Data and materials availability

All data will be available in some form to any researcher for purposes of reproducing or extending the analysis.

## Supplementary Materials

Figs. S1 to S3

## Materials and Methods

### Experimental Design

#### Subjects

Data were recorded from two female rhesus monkeys (Macaca mulatta, aged 7 and 8 years respectively, and weighing between 5 and 6 kg) renamed M1 and M2. Animal care, housing, and experimental procedures were in compliance with the National Institutes of Health’s Guide for the Care And Use of Laboratory Animals and approved by the Ethical board of Institut de Neurosciences de la Timone (ref 2016060618508941).

#### Alert monkey fMRI

The monkeys were first scanned for identifying Temporal Voice Areas (TVAs). All the details about the fMRI procedures are reported in Bodin et al. (2021) – in which the two monkeys were called M2 and M3. Here we give only a brief description of these details.

Functional scanning was done using an event-related paradigm with clustered-sparse acquisitions on a 3-Tesla MRI scanner (Prisma, Siemens Healthcare), equipped with an 8-channels surface coil (KU, Leuven). Ferrous oxide contrast agent (monocrystalline iron oxide nanoparticle, MION) was used for all the scanning sessions. Monkeys were trained to stay still in the scanner for a fixed period of 8 seconds to receive the reward. To avoid interference between sound stimulation and scanner noise, the scanner stopped acquisitions such that three repetitions of one of 96 stimuli (inter-stimulus interval of 250 ms) were played on a silent background. The 96 stimuli consisted in brief complex sounds from four main categories: human voices, macaque vocalizations, marmoset vocalizations and non-vocal sounds. Then, MION functional volumes were acquired using EPI sequences (multiband acceleration factor: 2, TR = 0.955 s). The analysis included 67 MION runs of M1 and 64 MION runs of M2. Voice selective areas were identified as those regions responding significantly more to conspecific (macaque) vocalizations versus non-vocal sounds (Fig. 1A).

#### fMRI-guided electrophysiology

Monkeys were chronically implanted with high-density microelectrode arrays (CerePort Utah Array, Blackrock Microsystems, Salt Lake City, UT, USA) to record extracellular activity in the fMRI-localized TVAs. Functional maps projected on the individual anatomical surfaces were used to calculate the exact position of the arrays. M1 was implanted with two 32-channels arrays in the anterior TVA (aTVA) of the right rostral Superior Temporal Gyrus (rSTG; parcellation from the D99 macaque brain template; Ts2 from the AC map macaque brain template) and one 32-channels array in the right frontal cortex. M2 was implanted with three 32-channel arrays in the left rSTG (Ts2), of which one in the aTVA. In this study, we analyzed only data collected from the arrays implanted in the aTVA (i.e., two arrays for M1 and one array for M2; Fig. 1A).

Electrical signals were amplified and processed using a RZ2 BioAmp Processor (Tucker-Davis Technologies, Alachua, FL, USA) and sampled at 24414 Hz. Raw data collected during recordings were high-pass filtered (300–5000 Hz), and spike sorting was performed offline using the fully automated algorithm MountainSort v5 (Chung et al., 2017) with the SpikeInterface package (Buccino et al., 2020). This algorithm detects, for each channel, individual clusters that are then filtered by applying specific thresholds to their quality parameters to identify single-neurons (Chung et al., 2017). Clusters with very low firing rates (FRs) (<0.5 Hz) were discarded. Finally, the mean waveforms of the remaining clusters were visually inspected to exclude neurons with irregular shapes.

To limit the possible inclusion of the same neuron across sessions, we selected sessions separated in time by at least 3 days (M1: 9 ± 4 (mean ± SD) days’ intervals between sessions on average; M2: 17 ± 8 days’ intervals on average). Our final dataset was composed of 475 single neurons, with 358 neurons from M1 across 11 recording sessions, and 117 neurons from M2 across 5 recording sessions.

#### Experimental setup and behavioral task

All recordings were performed in an acoustically insulated room. The monkeys sat in a primate chair with the head fixed by a non-invasive modular restriction mask (MRM) developed in our laboratory. Auditory stimuli were presented through a RZ6 Multi-I/O Processor (Tucker-Davis Technologies, Alachua, FL, USA) and transduced by two 8020 Genelec speakers, which were positioned at ear level 72 cm from the head and 60 degrees to the left and right. Stimuli were delivered at a sound pressure level of approximately 92 dB. Hand detection was achieved using two optical sensors.

Monkeys were trained to perform a pure tone detection task (Fig. 1B). This task was introduced to maintain the attention of the monkeys on the auditory stimulation. They were required to hold a bar with both hands for 1500-2000 ms to trigger the presentation of the sounds. In each trial, from three to seven stimuli (inter-stimulus interval: 280-540 ms) were played after which a 500 ms 1000-Hz pure tone was presented. The pure tone instructed the monkeys to release the bar to receive the juice reward (correct trials). If the monkeys released the bar before the pure tone presentation (false alarm trials) or did not release the bar (miss trials; upper reaction time: 250 ms), no reward was given.

#### Auditory stimuli

During each recording session, we presented multiple sets of stimuli with a fixed order. The stimuli within each set were presented in a randomized order, but all stimuli were presented once before any repetition occurred, ensuring a balanced number of repetitions. The stimulus sets were as follows:

“Voice identity”. The main set of stimuli in this experiment consisted of 25 natural vocalizations of rhesus macaques (kindly provided by Marc Hauser). Specifically, we included 5 “coo calls” from each of 5 individuals of macaque (duration time: 300-500 ms). The stimuli were collected from subjects that were unfamiliar with the experimental monkeys. Stimuli were resampled at 48828 Hz and normalized by root mean square amplitude. Finally, a 10 ms cosine ramp was applied to the onset and offset of the stimuli.

“Localizer”. At the beginning of each recording session, we presented a set of 12 macaque vocalizations and 12 non-vocal sounds. Macaque vocalizations (kindly provided by Marc Hauser) included different call types (coos, grunts, barks and screams). Non-vocal sounds included both natural and artificial sounds from previous studies from our group (Belin et al., 2000; Capilla et al., 2013) or kindly provided by Petkov et al. (2008) and Moerel et al. (2012). Stimuli were adjusted in duration so that all of them lasted 500 ms.

“Band-passed noise” (BPN). In each recording session, after the localizer, we assessed the tonotopic organization of the recorded areas, by means of five stimuli, each with a central frequency ranging from 125 Hz to 16000 Hz, spaced in 1.75 octave steps (band weight: ⅓ octave). The results regarding the tonotopic organization of the aTVA have been previously reported in Giamundo et al. (2024), where no clear tonotopic organization was observed in this region.

### Data Analysis

#### Spiking data preprocessing and neurons’ selection

Analyses were conducted using MATLAB (The MathWorks, Inc., Natick, MA, USA) and the open-source statistical software R. The dataset for this study comes from recording sessions in which each of the 25 voice identity stimuli was repeated at least 28 times in M1 sessions and 27 times in M2 sessions.

Spike times were saved at a resolution of 1ms. Peri-stimulus time histograms (PSTHs) were smoothed with a Gaussian kernel of 10 ms. All the data in this paper were aligned to the onset of sounds. Baseline activity was defined as the average FR during the 100 ms period preceding stimulus onset. To compute z-scores, the average response to each stimulus was normalized to standard-deviation units (SD) relative to the baseline. Since the duration of the macaque voice identity stimuli varied between 300 and 500 ms, we analyzed a 300 ms window following stimulus onset, corresponding to the minimum presentation time across all stimuli.

Auditory-responsive neurons were selected by calculating z-scored responses to each of the 25 voice identity stimuli in 10 ms bins throughout the stimulus presentation. Neurons were classified as auditory-responsive if their z-score exceeded 2 SDs for at least two consecutive bins in response to one or more stimuli (similar to methods from Perrodin et al., 2011; Giamundo et al., 2024). Only auditory-responsive neurons (76 from the most anterior array of M1; 97 from the most posterior array of M1; 44 from M2) were included in further analyses.

To compute the mean normalized responses of each neuron to the stimuli in Fig. 1E, we calculated the average response to each stimulus in a 200 ms window after onset, subtracted the baseline, and normalized by the absolute maximum across the stimuli.

#### Voice Selectivity Index

For each neuron, we calculated a Voice Selectivity Index (VSI) using the localizer stimulus set. The VSI was defined as:

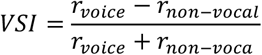

where *r*_*voice*_ is the average neuronal response to macaque vocalizations, subtracted of the baseline, in a 200 ms window following stimulus onset, and *r*_*non-vocal*_ is the average neuronal response to non-vocal sounds, subtracted of the baseline, in the same time window.

A VSI of 0 indicates equal responses to voices and non-vocal sounds. A VSI of 0.33 indicates a response twice as strong to vocalizations compared to non-vocal sounds; while a VSI of -0.33 reflects a response twice as strong to non-vocal sounds as to voices. These thresholds were used to classify neurons as voice-selective or non-vocal selective, consistent with previous studies (Perrodin et al., 2011; Giamundo et al., 2024). In cases where *mean*_*voice*_ > 0 and *mean*_*non-vocal*_ < 0, the VSI was set to 1; conversely, in cases where *mean*_voice_ < 0 and *mean*_*non-vocal*_ > 0, the VSI was set to -1 (Freiwald & Tsao, 2010; Giamundo et al., 2024).

#### Preference Index

To assess the preference of neurons for the different voice identities, we calculated for each neuron a preference index (PI) as:

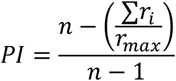

Where *n* is the number of identities, *r*_*i*_ the average neuronal response to identity*i*, subtracted of the minimum *r*_*i*_, in a 200 ms window following stimulus onset, and *r*_*max*_ the maximum *r*_*i*_. A value of 0 indicates an identical response to the five identities; a value of 1 indicates modulated discharge for only one identity (Umilta et al., 2007).

#### Decoding analysis

We used a maximum correlation coefficient classifier (MCC; as implemented in the MATLAB neural decoding toolbox (Meyers, 2013) to analyze the aTVA neuronal population or the different subpopulations of neurons (Voice-selective vs. Non-vocal selective; Identity-selective vs. Not identity-selective). The classifier was trained to discriminate between the five voice identities.

To train the classifier, trials (i.e., stimuli) were labeled based on their identity, and firing rates from trials and neurons were binned in 20 ms sliding windows, every 10 ms bins. We tested 44 consecutive bins, from -100 ms (i.e., a time window from -100 ms to -80 ms) to +330 ms from the stimulus onset. Note that these 20 ms bins are plotted such that the decoding accuracy is aligned to the center of each bin. For each bin, a different classifier was trained/tested.

Z-score normalization was applied to each neuron, to give equal weight to all the units regardless of firing rate. Then, the classifier was trained using a k cross-validation splits procedure, where k represents the maximum number of available trials for each condition for each neuron. Particularly, the classifier was trained using k – 1 splits and then tested on the remaining split. All possible train/test splits were tested and this process was repeated 50 times (i.e., 50 runs) with different subsets of trials. The classification accuracy from these runs was then averaged.

To assess whether the obtained decoding accuracies were above chance, we ran a permutation test that consisted of repeating the full decoding procedure 50 times with the labels of identities randomly shuffled. We obtained a null distribution of shuffled data, and the decoding results were considered significantly above chance if they were greater than all the shuffled data in the null distribution (p-value threshold of p = 1/ (50 * 44) = 0.0004). The latency of when the p-values are first above chance corresponds to the first time bin of 3 consecutive bins with p-values below the p-value threshold.

To determine the similarity between two classification accuracy distributions, we computed for each time bin a distribution-free overlapping index (η) using the overlapping package for R (Pastore & Calcagní, 2019). The overlapping index η represents the proportion of the overlapping area between the probability density functions of two distributions. In this sense, an overlapping index of η(A,B) = 0 indicates that f_A (X) and f_B (X) are distinct. Two distributions were considered as significantly different for η < 0.05 for at least 3 consecutive time bins.

For the identity-selective and non-identity-selective subpopulations, we also performed a cross-temporal decoding analysis to establish the temporal evolution of information coding as previously done (Meyers et al., 2008; Giamundo et al., 2022; Ceccarelli et al., 2023). This analysis results in a classification accuracy matrix where the values along the diagonal are calculated by performing training and testing on equivalent time bins. In contrast, different time bins for training and testing are used to calculate the off-diagonal values. We classified the stability of off-diagonal time points by implementing the method used in Ceccarelli et al. (2023).

#### Dimensionality reduction approach

We applied Principal Component Analysis (PCA) to reduce the dimensionality of the neuronal population’s responses. This method allowed us to represent responses to the 25 voice-identity stimuli as neural trajectories evolving in a low-dimensional state space.

For each stimulus and neuron, we computed the average spike density function (SDF) in 1 ms time bins, smoothed using a Gaussian kernel with a 10 ms width, across a time window from - 100 ms to +300 ms relative to stimulus onset. SDFs were further smoothed using 100 ms time windows. The resulting SDFs for all neurons and stimuli were organized into matrices. These matrices were concatenated while preserving the number of rows (corresponding to the number of neurons). The activity of each neuron was normalized by subtracting mean activity and dividing by SD. PCA was then applied to the concatenated matrix with dimensions N (neurons) × StTi (stimuli × time bins).

To examine how neural trajectories converged or diverged based on voice identity, we calculated the Euclidean distances between trajectories in the reduced-dimensional state space. Specifically, we averaged the pairwise distances (n=50) between trajectories corresponding to the same identity (Within IDs distance). Similarly, we averaged the pairwise distances (n=250) between trajectories from different identities (Between IDs distance). Statistical comparisons between the ‘Within IDs distance’ and ‘Between IDs distance’ were performed using bootstrapped two-sample t-tests (100,000 iterations, one-tailed), with Bonferroni correction applied for multiple comparisons, resulting in a corrected significance threshold of p=0.0001 (corresponding to p=0.05/401 time bins).

#### Representational Similarity Analysis

We performed representational similarity analysis (RSA) to compare neuronal data with acoustic data and evaluate theoretical models of voice identity representation.

In a first analysis, we generated time-resolved representational dissimilarity matrices (RDMs) from neuronal data. For each of the 25 voice identity stimuli, we pulled out a vector of z-scored averaged FRs from neurons in 10 ms time bins, using a ± 25 ms sliding window, from -100 to 300 ms relative to stimulus onset. At each time bin, a 25×25 RDM was generated by computing for each pair of stimuli the Euclidean distance between their population vectors. This resulted in a series of 41 Neuronal RDMs.

We compared these 41 Neuronal RDMs with a Model RDM representing a theoretical pattern of pairwise dissimilarities in our stimulus set corresponding to the ideal distinction between identities (referred as “voice discrimination” process) and to the ideal association of voices within each identity (referred as “voice recognition” process; Fig. 2A, right panel). Neuronal RDMs were also compared with an LTAS RDM consisting on the dissimilarity in long-term average spectrum (LTAS) for each pair of the 25 voice identity stimuli to account for differences in their low-level acoustical properties.

To compare the different RDMs, we applied two paradigms. In Paradigm 1, we computed time-resolved Spearman’s rank correlations (Bonferroni corrected p-value threshold of p = 0.05/41 = 0.0012) between the lower triangles (excluding the diagonal) of the ranked Neuronal RDMs and the Model RDM (i.e. 41 correlation values). In Paradigm 2, we computed the time-resolved Spearman’s rank correlations between ranked Neuronal and Model RDMs as in Paradigm 1, but controlling for the influence of LTAS by partialling out the LTAS RDM.

We further tested associations between Neuronal RDMs and Model RDM by comparing specific portions of ranked Neuronal RDMs corresponding to voice discrimination and voice recognition processes, predicted by the model. Comparisons were performed using bootstrapped two-sample t-tests (100,000 iterations, one-tailed; Bonferroni-corrected p-value threshold: p = 0.05/41 = 0.0012). The resulting voice discrimination and voice recognition curves were compared to baseline (-100 to 0 ms) using paired-sample t-tests with the same correction.

In a second analysis, we computed a single 25×25 Neuronal RDM over the 0-200 ms post-stimulus onset period, for five groups of identity-selective neurons and five groups of non-identity-selective neurons, defined by their preferred identity based on tuning profiles. This produced 10 Neuronal RDMs. These Neuronal RDMs were compared to five Model RDMs, each representing the ideal separation of one specific identity from all others. Planned comparisons involved analyzing Between IDs versus Within IDs portions of ranked Neuronal RDMs, predicted by the models (Fig. 3C, middle panels), using bootstrapped two-sample t-tests (100,000 iterations, one-tailed; Bonferroni-corrected p-value threshold: p = 0.05 / 50 = 0.001).

