## Supplementary figures for "Voice identity invariance by anterior temporal lobe neurons"

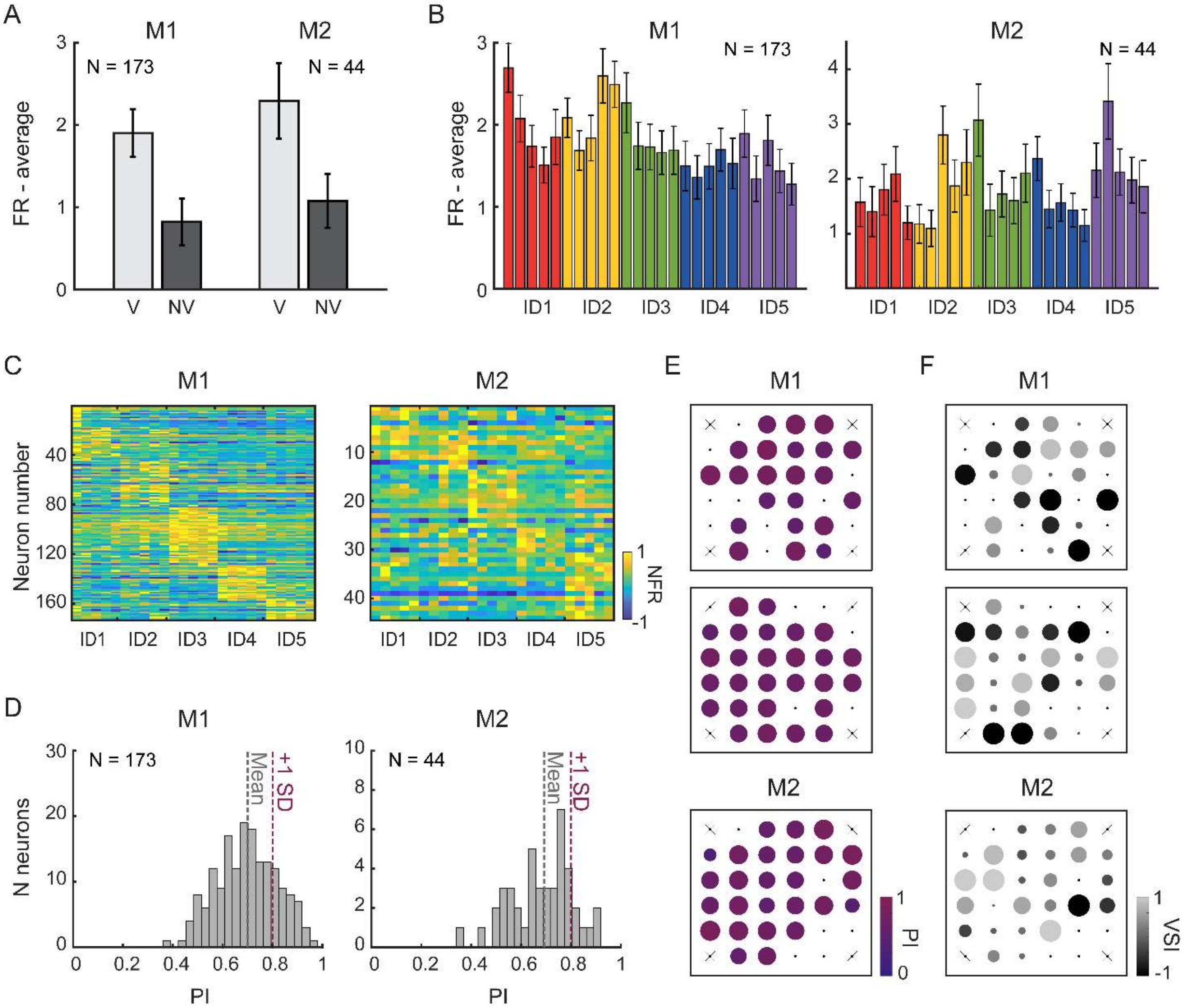


<u>Fig. S1. aTVA neurons responsiveness to voice identity in Monkey 1 (M1) and Monkey 2 (M2).</u> (A-B) Average population response (mean ± SE) to macaque vocalizations (V) and non-vocal sounds (NV) of the localizer stimulus set (A), and to the 25 voice identity stimuli (B), with the color corresponding to each identity. (C) Neurons responsiveness to the 25 voice identity stimuli. Each row represents the average response of a single neuron across the 200 ms post-stimulus onset. Neurons are sorted based on the identity eliciting their strongest response. (D) Histograms of Preference Indices (PIs), reflecting each neuron's selectivity for a single identity. A PI of 0 indicates equal responsiveness across all five identities, while a PI of 1 reflects exclusive responsiveness to a single identity. Vertical dashed lines represent the mean PI (Mean) and the mean plus one standard deviation (+1 SD). (E) Representation of Preference Indices (PIs) in electrophysiological recording sites. PIs were averaged across neurons recorded at each electrode of the arrays. (F) Representation of Voice Selectivity Indices (VSIs) contrasting macaque vocalizations vs. non-vocal sounds in electrophysiological recording sites. VSIs were averaged across neurons recorded at each electrode of the arrays. A VSI = 1 indicates full selectivity for macaque vocalizations, whereas a VSI = -1 indicates full selectivity for non-vocal sounds.

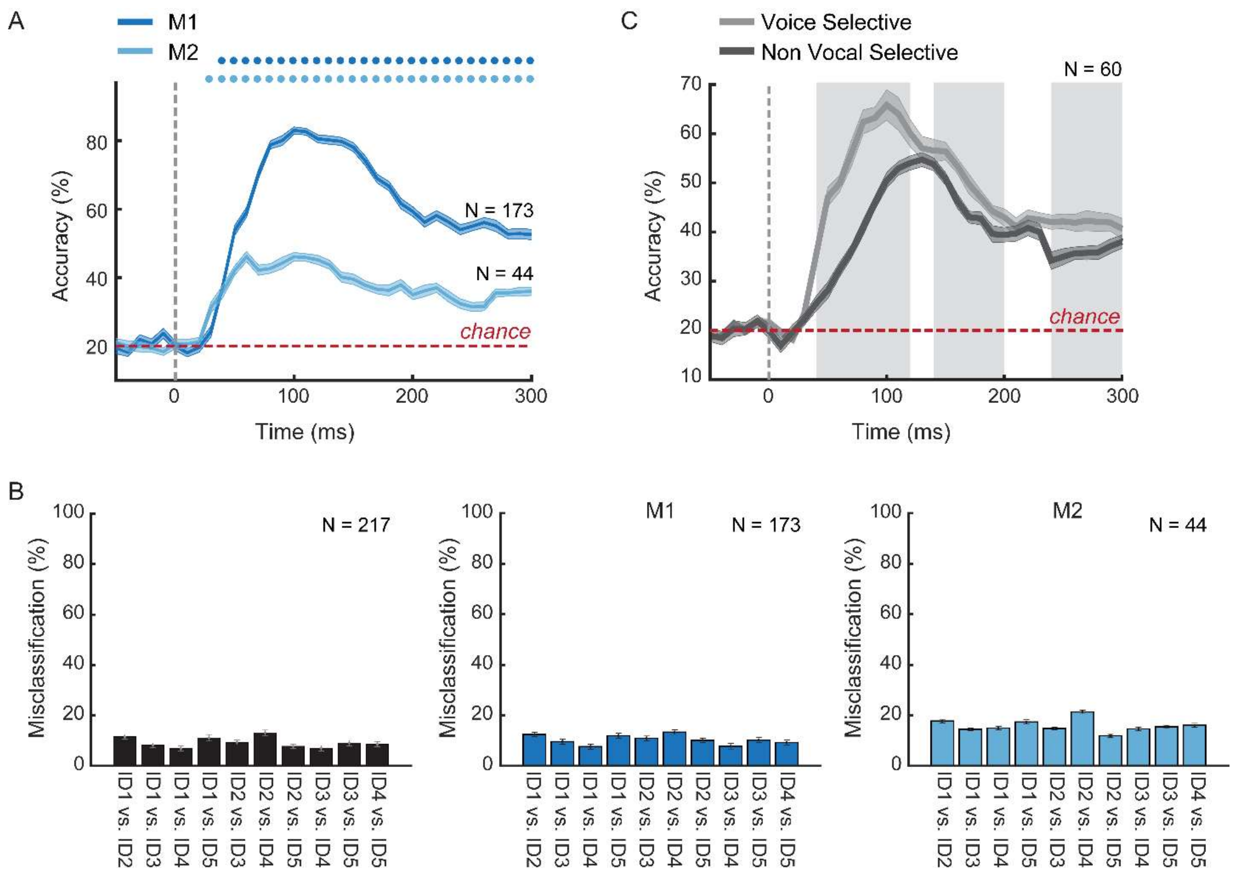


Fig. S2. Encoding of voice identity information at the population level. (A) Time-resolved classification accuracy (mean ± SE) of linear classifiers trained to discriminate between identities (chance level: 20%) based on the spiking activity of neurons in M1 and M2. Colored dots above the curves indicate time bins with significantly above-chance classification accuracy (permutation tests, $p < 0.0004$). (B) Misclassification rates for all identity pairs, computed using neurons from both monkeys (top panel), only M1 (middle panel) or only M2 (bottom panel). (C) Accuracy (mean ± SE) in the classification between identities (chance level: 20%) for voice-selective neurons (i.e. neurons with a VSI > 0.33) and non vocal-selective neurons (i.e. neurons with a VSI < -0.33). Voice-selective accuracy curve represents the average (± SE) classification accuracy from 40 subsamples of 60 neurons (on 92) that were randomly selected to have the same number of non vocal-selective neurons (N = 60). Colored dots indicate time bins with significantly above-chance classification accuracy (permutation tests, $p < 0.0004$). The gray shaded areas indicate time bins with significant difference between classification accuracies curves ($\eta < 0.05$).

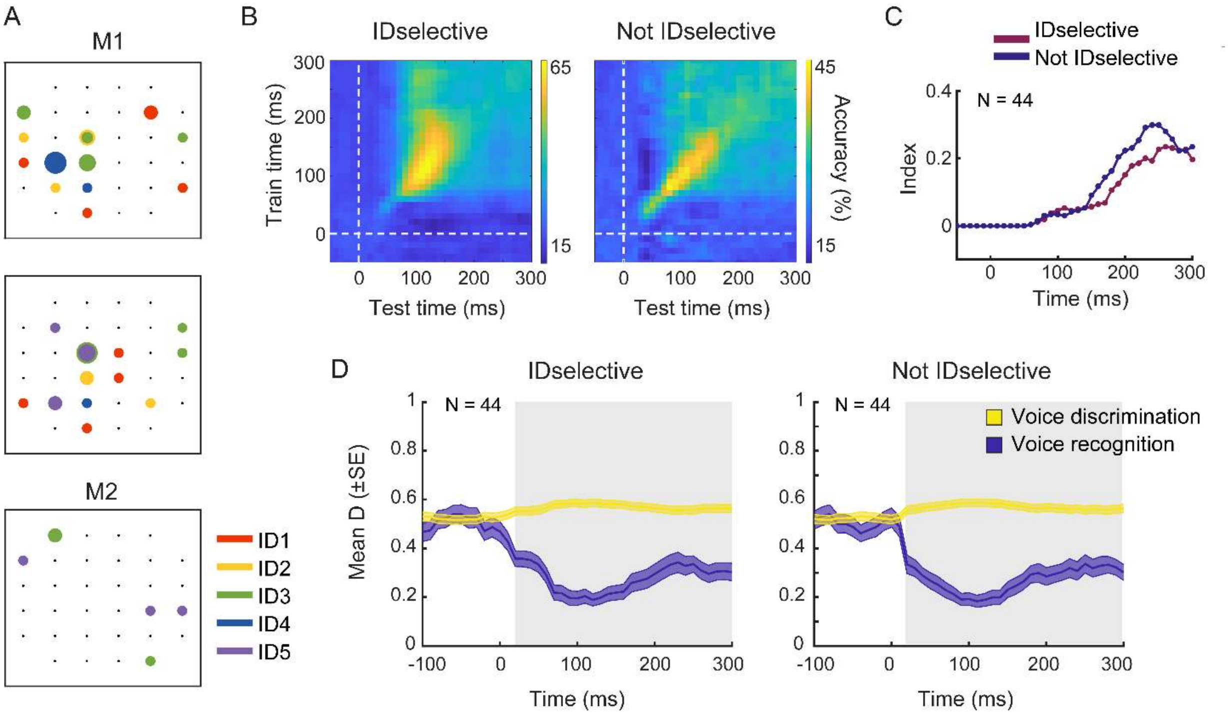


Fig. S3. Comparison of identity-selective versus non-identity-selective neurons. (A) Representation of N=44 identity-selective neurons in electrophysiological recording sites. Size of dots corresponds to the number of identity-selective neurons recorded from one site. The color code corresponds to the preference for one particular identity. (B-C) Cross-temporal decoding (B) and stability indices (C) for identity-selective and non-identity-selective subpopulations. (B) Cross-temporal decoding analysis evaluates whether the same neural code is employed at different time points for processing voice identity information, so examining the dynamic or static nature of population coding. In the plots, the y-axis represents the time bins used for training the classifier, while the x-axis represents the time bins used for testing. A dynamic coding pattern is indicated by a diagonal band of high accuracy (suggesting rapid decay of accuracy when testing at time points different from the training period). In contrast, static coding is characterized by broader, square-like regions of high accuracy, indicating consistent performance across time points. (C) Stability indices quantify the stability of the coding over time for each subpopulation. High stability index values indicate stronger temporal stability in neural coding. (D) Representation of neural patterns underling voice discrimination and voice recognition processes in both subpopulations, as done in Fig. 2D. The gray shaded area indicates time bins of statistically significant association with the theoretical model (bootstrapped two-sample t-tests, Bonferroni-corrected p = 0.0012).
